# Effects of cognitive load and years of experience on phase-amplitude coupling in simultaneous interpretation

**DOI:** 10.1101/2024.05.03.592346

**Authors:** Haruko Yagura, Hiroki Tanaka, Satoshi Nakamura

## Abstract

Simultaneous interpretation is a highly cognitively demanding task that requires constant attention switching between languages. Interest continues to grow in the contribution of phase-amplitude coupling (PAC), which involves the cooperative interaction of multiple oscillations and working memory. In this study, we established subjective definitions for cognitive load levels based on the subjective word familiarity of simultaneous interpretation, categorizing them as low, medium, or high. We then compared the changes in the PAC patterns between experienced interpreters and beginners. Experienced interpreters exhibited an increase in PAC, including theta-gamma PAC, which is linked to working memory, as well as theta-beta PAC, alpha-beta PAC and alpha-gamma PAC, with rising cognitive load levels in simultaneous interpreting. This suggests that experienced simultaneous interpreters choose a more adaptive neural processing strategy in response to the cognitive demands of interpretation language. In contrast, beginner interpreters do not show such changes in PACs, indicating either an underdeveloped or a different neurological approach to the cognitive load levels of interpretation language. The difference in PAC responses between the two groups reflects varying cognitive and interpretive strategies in the brain, where experienced interpreters might utilize more advanced neural mechanisms to manage higher levels of difficulty in simultaneous interpretation.

## Introduction

Simultaneous interpretation (SI) demands a very high level of multitasking ability. Simultaneous interpreters (interpreters) must start explicating new speech before they have finished interpreting the previous language.Interpreters require the crucial skill of quickly shifting their attention between different languages by delicately balancing listening and speaking while rapidly switching their focus between languages. Such ability for language-attention switching (language-switching) is developed through training and experience [1–3]. Furthermore, the brain function for language-switching improves with years of interpreting experience. However, there remains a lack of comprehensive neuroscientific understanding regarding how the brain responses of experienced interpreters dynamically change the brain’s response to language-switching for ongoing speech.

Cowan’s embedded-process model proposes that interpreters must concentrate on incoming speech, temporarily store its information in their working memory, and proceed with interpretation [4–7]. The model is comprised of three processes: long-term memory, activation, and focus of attention. Interpreting requires the simultaneous activation of extensive knowledge from long-term memory, which includes grammar and vocabulary, while focusing attention on working memory. Episodic memory, which is a type of long-term memory that draws from past experiences, plays a vital role in this process. By leveraging past experiences, interpreters are able to access and activate the necessary grammatical and lexical knowledge required for interpretation [8, 9]. Additionally, the prefrontal cortex is responsible for selecting information through the executive processes of working memory and attention by focusing on relevant information and activating it as needed [10–12]. Cowan’s embedded-process model significantly contributes to the understanding of language-switching in SI.

Previous studies demonstrated that with increasing SI experience, changes occur in brain activity that enhance the ability to switch attention. Several studies on the brains of interpreters have assessed the impact of interpreting experience on the brain from structural and functional aspects. In structural studies, Elmer et al. discovered that years of work experience in SI induces changes in the properties of the white and gray matter, and Becker et al. identified higher gray matter density in interpreters [13, 14]. Longitudinal studies by Hervais-Adelman et al. and van de Putte et al. concluded that the training of interpreters influences the thickness and connectivity of their brain regions [15, 16]. In functional studies, Rinne et al. observed activation in brain regions associated with language-attention switching, and Elmer et al. noted activation in specific areas during interpretation in the second language and in different interpretation directions [17, 18]. Hervais-Adelman et al. discovered the involvement of the caudate nucleus and the ventral striatum in controlling language output [19]. EEG studies highlight functions involved in language-switching during SI. Grabner et al. observed an increase in the theta and alpha bands, and Elmer et al. saw an amplification of the N400 response in reverse-direction tasks [13, 20, 21]. Klein et al. found excessive interhemispheric connectivity in the alpha band, and Koshkin et al. noted that task-irrelevant sounds during SI causes changes in brain responses, indicating increased working memory load [22, 23]. Yagura et al. explored the changes in 40-Hz auditory steady-state responses while interpreters with varying lengths of SI experience listened to a 60-second audio source. The study confirmed that phase synchronization was influenced by the number of years of SI experience [24, 25].

It has recently been reported that neural oscillations do not function independently, but rather collaborate with one another to maintain working memory. Cross-frequency coupling occurs when interactions take place between different frequency bands in brain oscillations. There are four forms of Cross-frequency coupling: phase-amplitude, power-power, phase-phase, and phase-frequency coupling [26–28]. Among these, phase-amplitude coupling (PAC) is a prevalent form of Cross-frequency coupling, where the amplitude of high-frequency oscillations is modulated by the phase of low-frequency oscillations [29, 30]. PAC is characterized by the interaction between the phases of low-frequency oscillations, typically in the theta band, and the amplitude of high-frequency oscillations, usually in the gamma band [29, 31]. In PAC, a phase, which indicates the repeating position of the oscillations, influences the amplitude, which represents the strength of the oscillations [29, 31, 32]. Furthermore, the role of PAC in working memory is involved in the integration of various neuronal oscillations.

Recent research has revealed a significant relationship between theta-gamma PAC and information processing in working memory, as demonstrated by various studies [33–36]. Both theta and gamma bands have separate functions in cognitive processing. Theta bands, mainly in the hippocampus, help with learning and memory formation and are involved in incorporating new information and combining memories for long-term storage [37, 38]. Gamma bands, on the other hand, are high-frequency brain bands that integrate information across the brain and process complex tasks [39, 40]. They are crucial for higher brain functions like consciousness, perception, attention, and problem-solving. The combination of theta and gamma bands is especially critical for tasks requiring immediate language processing, sustained attention, information retention, and updating the processing of information in working memory [33–35]. This aspect includes regulating the timing and synchronization of neural activity, facilitating transitions between distinct information-processing tasks, encoding and retrieving memories, updating language, engaging in predictive processing, and organizing memory [33]. In tasks related to information organization, theta-gamma PAC plays a crucial role by enabling specific information to be activated or reactivated at specific times for facilitating information-processing adjustments [33, 36, 41]. A theta-gamma PAC is involved in attention-selection mechanisms, which reduce interference from other information sources and support a focus on specific information [42]. On the other hand, as the cognitive load of working memory increases, the theta-gamma PAC also increases, supporting stimulus processing and conveying information about perceptual and mnemonic representations [43]. Several studies have consistently reported an increase in the theta-gamma PAC, particularly in the frontal brain regions, as cognitive demands escalate [11, 44, 45]. Intracranial electroencephalography (iEEG) studies have further confirmed that theta-gamma PAC intensifies with heightened cognitive load, particularly in the hippocampal region [43, 46]. In such working memory tasks as the delayed match-to-sample visual working memory task, partial occipital cortices have been implicated in the theta-gamma PAC [47]. Therefore, it plays a vital role in coordinating various processes within the brain and enhancing information processing, memory, attention, and cognitive functions.

The alpha-gamma PAC plays a crucial role in the coordination of memory and attention [35, 48–50]. Alpha oscillations are particularly important in processes related to attention and memory because they help process information by suppressing irrelevant details and focusing on task-related information [35, 51, 52]. Gamma oscillations synchronize information within the brain, supporting efficient information processing and the maintenance and recall of essential data in the working memory [41, 53]. The significance of alpha-gamma coupling lies in its impact on the changes in brainwave activity linked to the working memory content and the nature of the task. Shifts in working memory content are accompanied by corresponding changes in brainwave activity. For instance, when working memory content moves from multiple items to specific visual or spatial information, theta activity tends to shift to alpha activity [54]. During the delay period of working memory, theta oscillations are more frequent in tasks requiring the encoding of sequential multiple working memory items; alpha oscillations are typically observed in tasks involving the simultaneous retention of visual or spatial information [35]. Similarly, pictorial recognition tasks, analyzed using EEG data, have revealed activation patterns in the prefrontal-parietal regions for the alpha-gamma PAC [55]. Verbal-delayed-match-to-sample tasks have linked gamma activity in the frontal regions to the alpha-gamma PAC [56]. In visual-delayed matching-to-sample tasks with varying working memory loads, both the frontal and posterior cortices have exhibited increased alpha-gamma PAC activity [57]. These findings highlight the importance of alpha-gamma coupling in understanding memory, attention, and cognitive processes, especially regarding how brainwave activity is influenced by the working memory content and the nature of the task.

The PAC of the beta bands is a critical factor in performing cognitive tasks, particularly working memory and visual match-to-sample tasks. Research by Daume et al. revealed that during the working memory delay period, increased theta-bata PAC in the medial temporal lobe is linked to more efficient cognitive processing [58]. The beta-band PAC helps synchronize different brain regions. For instance, Daume et al. observed phase synchronization in the beta band between the medial temporal lobe and the temporo-occipital areas [59]. Dimitriadis et al. concluded that the strength of a beta-band PAC decreases as the complexity of the arithmetic tasks increases, suggesting its role in adapting information processing based on task difficulty [60]. Additionally, a decrease was reported in the alpha-beta PAC in the occipital lobe during the encoding of episodic memories [61]. Recent studies have also highlighted the heightened connectivity between the alpha-beta and gamma bands within the dorsal and ventral visual pathways, particularly during the more demanding phases of N-back tasks [57]. Conversely, an increase in the theta-beta PAC was observed in the left inferior temporal cortex during visual working memory tasks, a region associated with visual object memory [59]. These findings illustrate that the beta-band PAC is crucial for supporting the efficient functioning of the brain, particularly in maintaining working memory and processing complex cognitive tasks. However, it remains unclear how these PACs work during SI, which is multitasking and intensely demanding.

Tong et al. pilot study’s examined the application of PAC in assessing the cognitive load associated with interpretion from the first language (native language: L1) to the second language (L2) [62]. L1-L2 language-switching during high interference results in a larger alpha-beta PAC for bilinguals, suggesting that PAC rises with higher cognitive load. Since this study focused on weak bilinguals with rather low language proficiency, the internal factors among the participants were not uniform. On the other hand, Lizarazu et al. explored the relationship between PAC and language proficiency comprehension [63] and examined the brain activity in native Spanish speakers who listened to reversed audio (harder to understand) and normal direction audio (easier to understand). Their results showed that theta-gamma PAC’s strength is linked to language comprehension. Higher Basque language proficiency (non-native language) was associated with stronger coupling; no such difference was observed for Spanish (native language). This result shows that the theta-gamma PAC increases with language comprehensibility and proficiency level.

However, these studies indicate that it remains unclear how much both the increase in cognitive load and the level of language proficiency affect PAC. Nor do they provide detailed information about the relationship with other types of PACs. Various studies have focused on working memory, where participants were assigned specific tasks to concentrate their attention. On the other hand, interpreters must simultaneously comprehend spoken language, translate it, and offer prompt and precise translations for subsequent discourse. They must concurrently perform numerous tasks, a task that requires a higher level of cognitive adaptability than simple cognitive operations. Such sophisticated multitasking places a considerable burden on interpreters, necessitating their ability to deliver high-quality performance and swift responses [4, 7], even though no investigations have delved into PACs related to SI. The main goal of our study was to investigate whether the cognitive load associated with interpreting during SI leads to an increase in PAC. We also anticipate that these responses will become more pronounced as interpreters gain experience. The following are the primary objectives of our research:

1. Investigating the relationship between SI’s cognitive load and PAC increases
2. Investigating the relationship between years of SI experience and PAC increases We analyzed the PACs using the data collected by Yagura et al. [24, 25].

## Materials and Methods

### Participants

Twenty two female Japanese interpreters were divided into groups of experienced (seven with at least 15 years of SI experience, age: mean = 56, SD = 8.8) and beginners (15 with at least one year of SI experience, age: mean = 51.2, SD = 4.3)(experienced or beginners). A Welch’s t-test showed no difference in age between the two groups: age: *t*_(21)_ = 1.5, p = 0.16, E-group: n = 7, mean = 56.71, SD = 8.8, B-group: n = 15, mean = 51, SD = 4.3).The number of experienced simultaneous interpreters capable of Japanese to English simultaneous interpretation is limited. A translation company selected all the participants and grouped them into beginner and expert groups based on their own criteria. The differences in SI difficulty by years of SI experience were assessed through objective evaluations [24, 25]. All subjects had undergone identical SI training, including shadowing (SH), and had approximately three years of experience in consecutive translation. The EEG data was collected between February and July 2019. All of the data collection processes were approved by ethical committees at the Nara Institute of Science and Technology. At the beginning of the EEG recording, we explained the procedure to the participants, obtained their written informed consent, and ensured this was documented in writing.

### Task sequence

This study conducted experiments to examine the cognitive load during simultaneous interpretation (SI) from Japanese to English and shadowing (SH) in Japanese. The SI task involved interpreting Japanese radio news in English, whereas the SH task involved listening to and repeating the same Japanese radio news. The order of task presentation is shown in Fig 1. Both tasks utilized the same audio material, with the SI task always preceding the SH task, to prevent cognitive load reduction due to content repetition. This study focused on using unedited, natural audio material to replicate the cognitive load experienced in real interpreting environments. Participants listened to eight randomly presented 60-second segments of Japanese radio news, which were neutral in emotional impact and rich in colloquial expressions, to minimize individual differences due to non-verbal impressions. To examine the impact of word familiarity on cognitive load, we intentionally withheld information about the content of the interpretation prior to the study. This approach allowed us to assess the cognitive load experienced by all participants in response to unfamiliar words. For more information on this task sequence, please refer to previous studies [24, 25].

**Fig 1.**
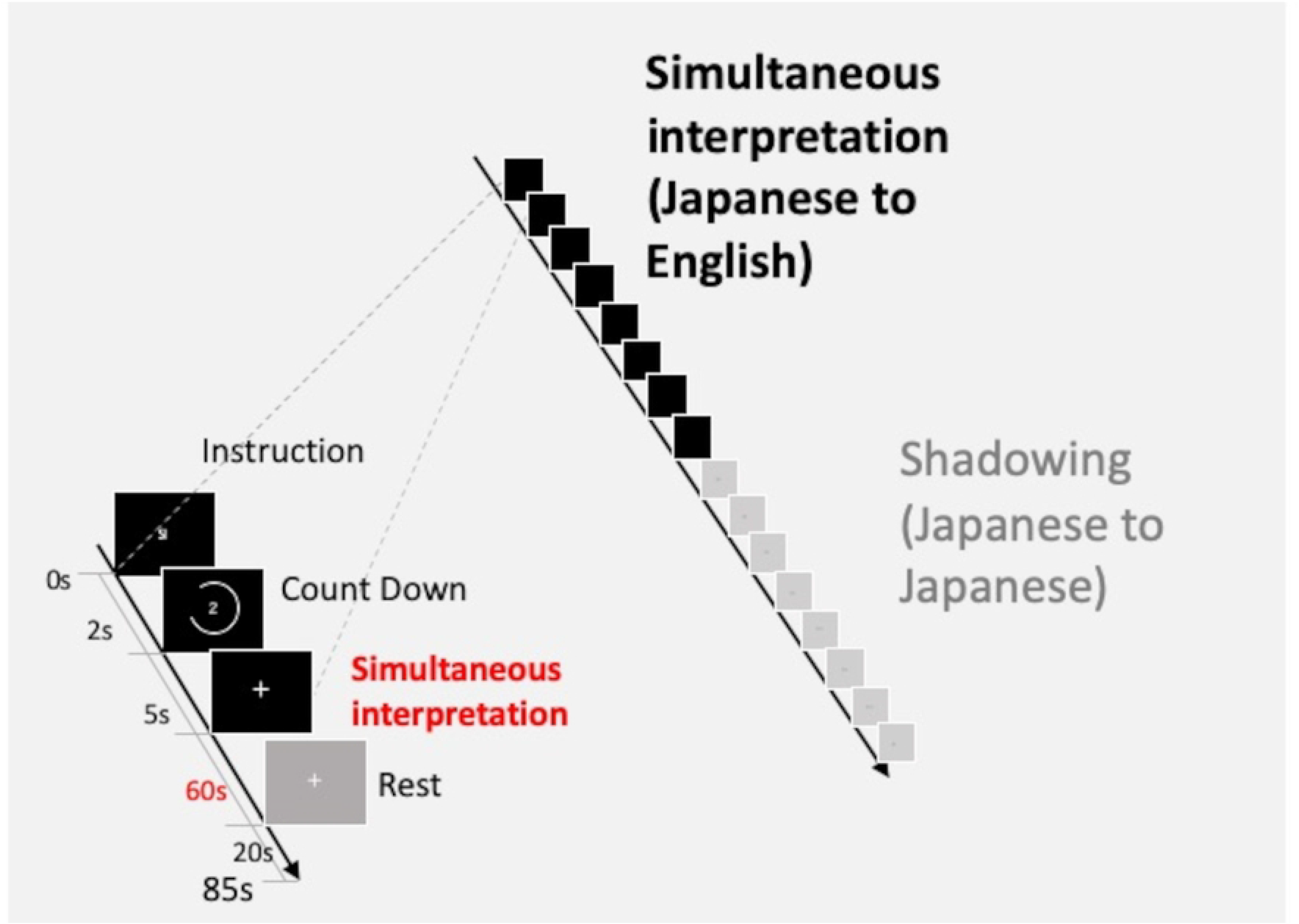
The task sequence during EEG measurement in simultaneous interpretation. This study focused on measuring cognitive load during simultaneous interpretation. The figure presented below illustrates a revised version of the task diagram, adapted to fit the objectives of this study, that has been previously reported in prior research [24].

### EEG data acquisition and prepossessing

The EEG signals were recorded by a Cognionics Quick-30 Dry EEG headset with 29 electrodes (excluding one for the reference electrode). We utilized a finite impulse response bandpass filter to process the recorded signals, which exhibited a passband ranging from 0.1 to 100 Hz and a sampling rate of 500 Hz, implemented through the MNE-Python [64]. We eliminated the eye movements and blinks using independent component analysis. This study explored the general relationships within the brain by focusing on PAC activity and the complexity of tasks like SI. PACs are frequently observed throughout distant parts of the brain [57, 61, 65]. To better understand the comprehensive relationships and functions of PAC across the entire brain, we performed analyses using all 29 channels without limiting ourselves to specific areas of interest.

### Sound stimuli

We used eight, 60-second Japanese radio news segments for the SI. The auditory repetitive stimuli were 40-Hz clicks (sampling rate: 8192-Hz), consisting of a 10-ms pulse width repeated every 25 ms. To subjectively evaluate the differences in the difficulty of interpretion that depends on the years of SI experience, the results showed that beginner SIs felt more difficulty during SI tasks than experienced SIs [24, 25].

### Subject evaluations to interpretation load

After the EEG recordings, all participants were presented with five questions related to each of the eight radio news segments. These questions pertained to the SI difficulty: Q1: interpretation achievement, Q2: topic familiarity, Q3: voice speed, Q4: the ease of hearing the voice, and Q5: overall interpretation difficulty. Responses to these questions were evaluated on a 5-point rating scale (1: easiest, 5: most difficult). Participants were not provided with interpretation materials in advance; instead, these materials were given to them just before the EEG experiment to ensure that differences in prior vocabulary knowledge did not introduce variations in cognitive load. Specifically, we focused on Question 2, “How familiar are you with this topic?” Here participants evaluated the familiarity of words on a scale from 1 to 5 (1: very familiar, 5: unfamiliar). The responses to Question 2 from all the participants (related to the eight news segments) were sorted in ascending order. Subsequently, these ratings were categorized into three difficulty levels (low, medium, and high), ensuring that each category contained basically the same number of responses (low: 59, medium: 58, and high: 59).

1. Cognitive load is low: very familiar;
2. Cognitive load is medium: medium familiarity;
3. Cognitive load is high: unfamiliar.

### Compute phase-amplitude coupling

We delineated specific frequency bands for conducting PAC analysis. The lower-frequency band, comprised of the theta and alpha ranges, was set from 3- to 13-Hz; the higher-frequency band, linked to amplitude modulation, encompassed the beta and gamma ranges from 14- to 55-Hz. We investigated the PAC properties by implementing a methodological approach that encompassed two primary stages. The initial stage involved time-frequency decomposition, which was achieved by applying the oscillation transform method to extract both the phase and amplitude information from the neural data under investigation. This approach enabled effective understanding of the temporal dynamics of oscillatory neural activity.

The second step involved surrogate data analysis to ascertain the statistical significance of the observed PAC. We achieved this by employing surrogate mean subtraction techniques to discern true PACs from spurious correlations within the data. The calculation process encompassed four essential steps. Initially, we extracted phase and amplitude information from the predefined frequency bands, as previously mentioned. Subsequently, we quantified the coupling strength between the phase of the lower-frequency oscillations and the amplitude of the higher-frequency oscillations. We ensured the reliability of our results by normalizing the strength of the PAC to mitigate potential spurious correlations. In our research, we categorized the PACs based on specific frequency ranges for a low-frequency phase (pha) and a high-frequency amplitude (amp). The following are the PAC categories: theta-beta PAC (pha: 0-4 Hz, amp: 13-27 Hz), theta-gamma PAC (pha: 0-4 Hz, amp: 28-40 Hz), alpha-beta PAC (pha: 5-9 Hz, amp: 13-27 Hz), and alpha-gamma PAC (pha: 5-9 Hz, amp: 28-40 Hz). Based on prior research by Lizarazu et al., the range for the high-amplitude gamma-band was set to 27-40 Hz [63]. We used Python’s Tensorpac (version 0.6.5) to calculate the PACs [66].

### Modulation index

The modulation index (MI), which is an indicator used in the analysis to quantify PAC, is known for its ability to provide reliable measurements in noisy environments and measurements independent of the signal’s amplitude [32]. PAC’s existence is characterized by the deviation of amplitude distribution *P* from a uniform distribution in the phase-amplitude plot. MI is an application of the Kullback-Leibler distance, which is widely used in statistics and information theory to estimate the difference between two distributions [67]. Thus, MI is a constant Kullback-Leibler distance of *P* from a uniform distribution. Expressed in the equation, the Kullback-Leibler distance of discrete distribution *P* and distribution *Q* is defined as (1):

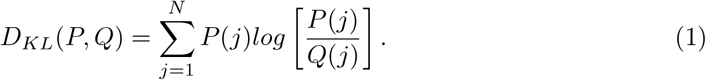

Note that the Kullback-Leibler distance formula resembles the definition of the Shannon entropy (*H*) of distribution *P*, given by (2)

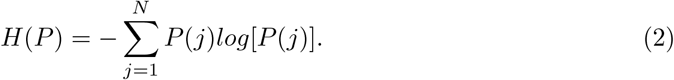

In fact, the Kullback-Leibler distance is related to the Shannon entropy by the following formula (3):

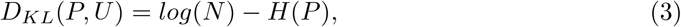

where *U* is the uniform distribution. Note further that *log*(*N*) is the maximal possible entropy value, which happens precisely for the uniform distribution. MIs are defined by dividing the Kullback-Leibler distance of observed amplitude distribution (*P*) from the uniform distribution (*U*) by Eq. (4):

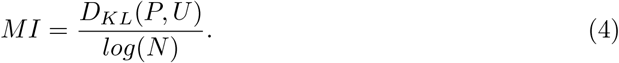

To verify whether the obtained original MI is statistically significant, a null distribution of MI was created by shuffling the trial and phase information assuming no coupling (200 permutations) at a sampling rate of 256 Hz to verify whether the distribution of the original MI is significant compared to a null distribution of the MI. We generated a null MI with the swap-amplitude time-block algorithm [32].

### Overview of MI statistical analysis flow

In our research, we examined the mean MI values for each type of PAC across all the channels to explore the impact of the load level and the participants’ work experience on these MI values. To compare the MI values across various load levels and work experience groups, we utilized the Kruskal-Wallis rank sum test and the Dunn’s post-hoc test. Our analysis elucidated the effects of cognitive load and work experience on PAC by categorizing the average MI values into three cognitive load levels (low, medium, and high) and two work experience groups (experienced and beginner). We focused on several PAC types: theta-beta PAC, theta-gamma PAC, alpha-beta PAC, and alpha-gamma PAC [62, 63, 65]. Additionally, we calculated the correlation coefficients to evaluate the relationship between the PACs and the cognitive load levels. Finally, we created a topography map using the MI values to visually represent the PAC changes. Our analysis quantified the relationship between the PACs and various cognitive load levels, clarifying the relationship between neural dynamics and workload during SI.

## Results

### Subjective evaluation analysis of the difficulty of interpretation

We used 2-way variance analyses (ANOVA) to examine the effects of work experience (experienced and beginner interpreters) and cognitive load levels (low, medium, and high) on these data. The results of the post-hoc tests following the ANOVA are shown in Fig 2 (a) and (b). Initial analysis revealed that the data were not normally distributed, as indicated by the results of the Shapiro-Wilk test (*p <* 0.001). Thus, we applied an aligned rank transform 2-way ANOVA for a non-parametric data-set. Significant main effects were observed in the subjective evaluation values across different cognitive load levels (cognitive load levels, *F* = 11.8, *p <* 0.001, 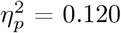: work experience, *F* = 12.0, *p <* 0.001, 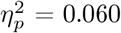). The interaction effect between work experience and cognitive load levels was not statistically significant, suggesting that their effects on subjective evaluation are independent of each other (interaction of cognitive load levels and work experience: *F* = 1.25, *p* = 0.287, 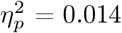).

**Fig 2.**
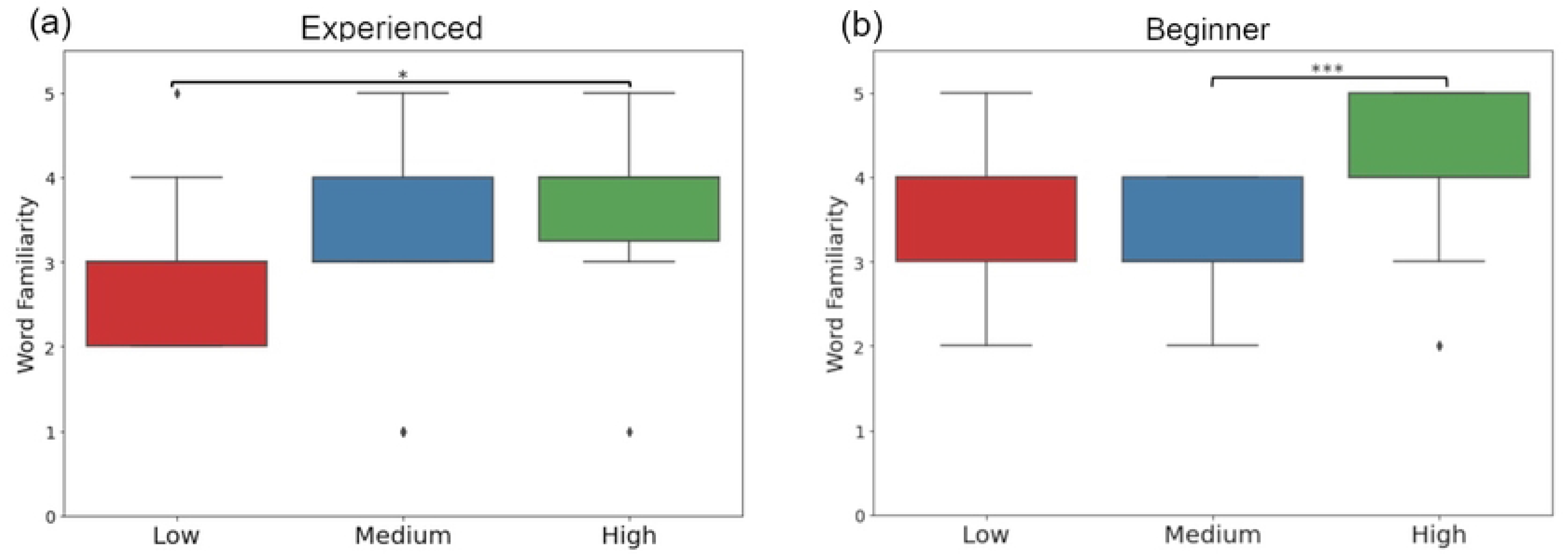
Subjective evaluations on interpreter difficulty during simultaneous interpretation across different years of interpreter experience. (a) Experienced interpreters. (b) Beginner interpreters. X-axis: levels of interpreting difficulty (Low, Medium, High). Y-axis: ratings of familiarity with the interpreted content on a scale from 1 to 5, where 1 indicates very familiar and 5 indicates unfamiliar. *p-values*: *p <* 0.05^*^, *p <*0.001^***^

Additionally, post-hoc tests using the Kruskal-Wallis test revealed significant differences in cognitive load levels within both the experienced and beginner groups (Experienced interpreters: *H* = 8.86, *p* = 0.011, Beginner interpreters: *H* = 18.19, *p <* 0.001). Following this, a Dunn’s post-hoc test with Bonferroni correction further emphasized these differences. Specifically, within the experienced group, a significant difference was observed between the high and low levels (experienced: high *>* low, *p* = 0.012) (Fig 2(a)). Conversely, in the beginner group, the difference was most pronounced between the high and medium levels (beginner: high *>* medium, *p <* 0.001)(Fig 2(b)).

To examine the overall difference in interpreter experience, we compared the subjective evaluation scores related to word familiarity between the experienced and beginner interpreters. A Mann-Whitney U test underscored a significant disparity in the subjective evaluation values, where beginners scored higher than their experienced counterparts (*U* = 2519.0, *p* = 0.0046, experienced: mean = 3.23, SD = 1.08, beginner: mean = 3.73, SD = 0.83).

According to the results of our subjective evaluation, we expect that beginner interpreters will perceive a greater cognitive load during the interpreting process than experienced interpreters. Furthermore, our results will show that the length of the work experience and cognitive load levels are performed independently of each other.

### Two-way ANOVA and correlation analysis between cognitive load levels and MI across PAC types

We conducted a 2-way ANOVA to examine the interaction between SI experiences as experienced and beginner groups and the cognitive load levels in three levels as low, medium, and high on the MI, which represents the PAC strength. Our analysis involved six types of PACs (Table 1). We conducted an aligned rank transform 2-way ANOVA because the data-set did not follow a normal distribution (Shapiro-Wilk test: *p<*0.05). The ANOVA results in Table 1 show that work experience plays a significant role in every PAC (*p <* 0.05). Especially, the interaction between work experience and cognitive load levels showed high significance across all PACs. In addition, it reveals the presence of notably high F-values and partial eta-squared 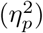 for the theta-beta and alpha-beta PACs (Table 1). Based on these considerations, SI seems characterized by a multitude of oscillation interactions. In other words, it is not solely dependent on a single PAC, but rather on a comprehensive response that is sustained by a variety of brain oscillations. Specifically, we expect that PAC’s influence on cognitive load levels will increase with an accumulation of experience.

**Table 1.** Interaction of cognitive load labels in SI and work experience on 2-way ANOVA. *p-values*: ^***^*p <* 0.001,^**^*p <* 0.01 and ^*^*p <* 0.05

| PAC types | Load |  |  | Work experience |  |  | Load vs Work experience |  |  |
| --- | --- | --- | --- | --- | --- | --- | --- | --- | --- |
| | *F-value* | $\eta_p^2$ | *p-value* | *F-value* | $\eta_p^2$ | *p-value* | *F-value* | $\eta_p^2$ | *p-value* |
| Theta-beta PAC | 2.18 | 0.020 | 0.038* | 19.13 | 0.089 | *** | 14.2 | 0.513 | *** |
| Theta-gamma PAC | 4.44 | 0.047 | 0.006** | 2.34 | 0.012 | *** | 7.37 | 0.484 | *** |
| Alpha-beta PAC | 2.93 | 0.027 | 0.017* | 20.15 | 0.093 | *** | 14.9 | 0.500 | *** |
| Alpha-gamma PAC | 3.71 | 0.038 | 0.005** | 10.16 | 0.052 | *** | 8.14 | 0.461 | *** |

Furthermore, Table 2 presents the correlations between work experience and cognitive load levels for each PAC. In the experienced group, a moderate positive correlation ranging from about 0.4 to 0.5 was observed between the cognitive load levels and MI. In contrast, the beginner group showed a negative correlation ranging from about -0.1 to -0.2. The effect size was similar for the experienced and the beginners. The results imply that work experience has different effects on PAC, suggesting that experienced participants and beginners use their brains differently when handling the cognitive load levels on SI. These analysis results validated our hypothesis that an increase in cognitive load during SI leads to an increase in PAC, and that this effect is influenced by the interpreter’s years of experience in SI.

**Table 2.** Correlation results of relationship between work experience and different PAC types. *p-values*:^***^*p <* 0.001 and ^*^*p <* 0.05

| Work experience | PAC types | *r* | Spearman's *p-value* | *cohend's d* |
| --- | --- | --- | --- | --- |
| Experienced | Theta-Beta PAC | 0.422 | *** | 1.65 |
| Experienced | Theta-gamma PAC | 0.426 | *** | 1.66 |
| Experienced | Alpha-beta PAC | 0.447 | *** | 1.71 |
| Experienced | Alpha-gamma PAC | 0.434 | *** | 1.65 |
| Beginner | Theta-beta PAC | -0.219 | 0.044* | 1.65 |
| Beginner | Theta-gamma PAC | -0.124 | 0.259 | 1.66 |
| Beginner | Alpha-beta PAC | -0.206 | 0.059 | 1.71 |
| Beginner | Alpha-gamma PAC | -0.111 | 0.261 | 1.68 |

### PAC changes on topography with more cognitive load by interpretation experience

Cognitive load effect on PAC dynamics is depicted in Fig 4, which show disparities in cognitive load levels (high-medium, medium-low, and high-low) between groups of experienced and beginner interpreters, as determined by average MI values for each of the 29 examined channels. The difference between the MI values of experienced and beginner interpreters under increasing cognitive load is clearly depicted in Fig 4, where experienced interpreters exhibit an increase in MI values across a variety of brain regions, such as the frontal, occipital, and temporal regions. This result indicates that multiple areas of the brain are engaged in response to cognitive load in PAC. In beginner interpreters, an increase in MI values was observed only in the parietal region under low cognitive load conditions (Fig 4 (a)-(d):B). As the cognitive load increased, this response disappeared, suggesting that an increase in cognitive load affects the distribution of brain activity in beginner interpreters. In contrast, experienced interpreters seem to effectively engage their entire brain in response to an increased cognitive load. We hypothesize that the strategies of brain utilization in response to cognitive load differ significantly between beginner and experienced interpreters (Figs 4 (a)-(d):E). Furthermore, Fig 4(b) shows that experienced interpreters have lower MI values in the alpha-beta PAC, suggesting a unique pattern in their brain activity [68].

**Fig 3.**
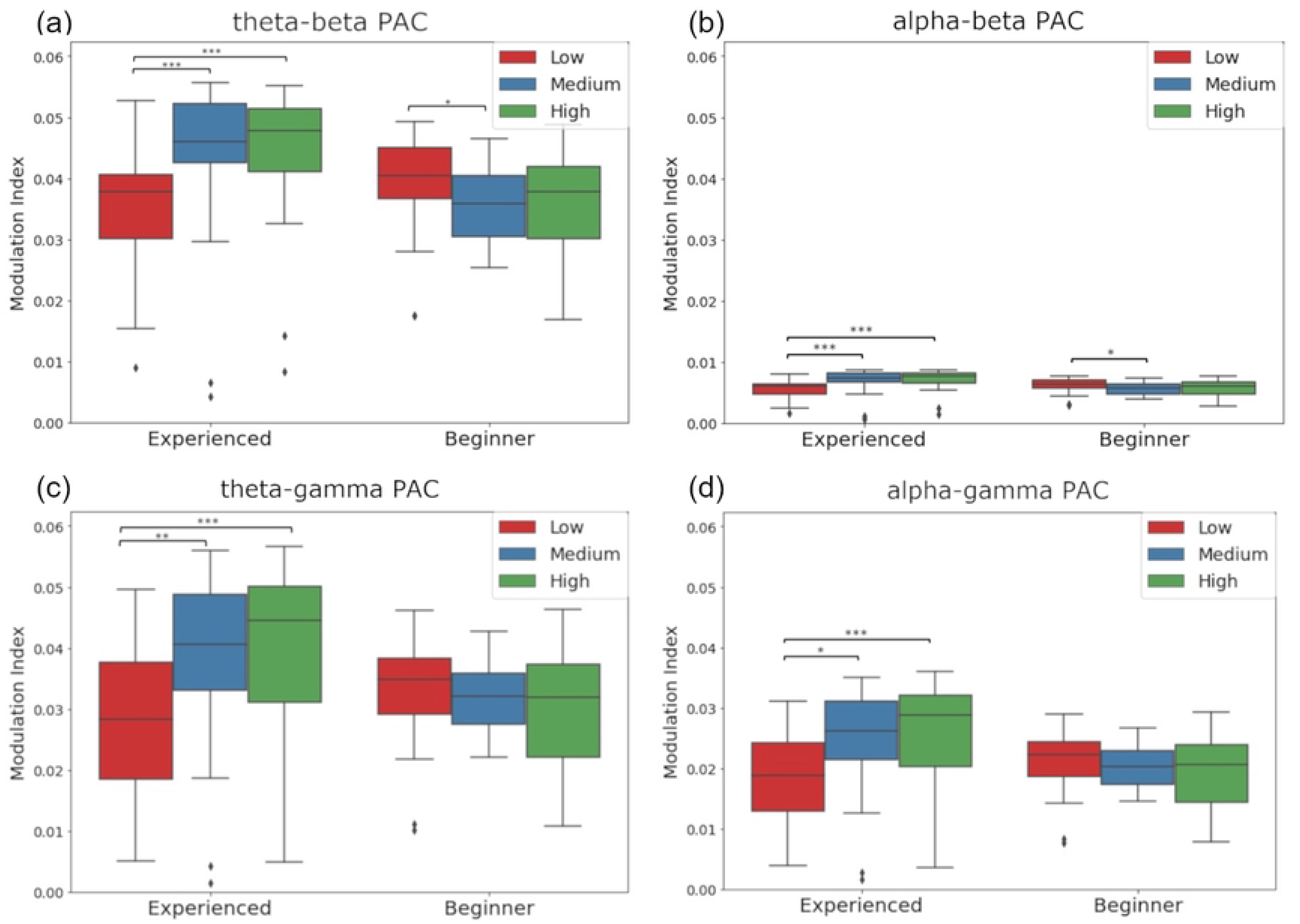
Dunn’s post-hoc test of cognitive load levels within each PAC. (a) theta-beta PAC. (b) alpha-beta PAC. (c) theta-gamma PAC. (d) alpha-gamma PAC. *p-values*: *p <* 0.05^*^, *p <* 0.01^**^, *p <*0.001^***^

**Fig 4.**
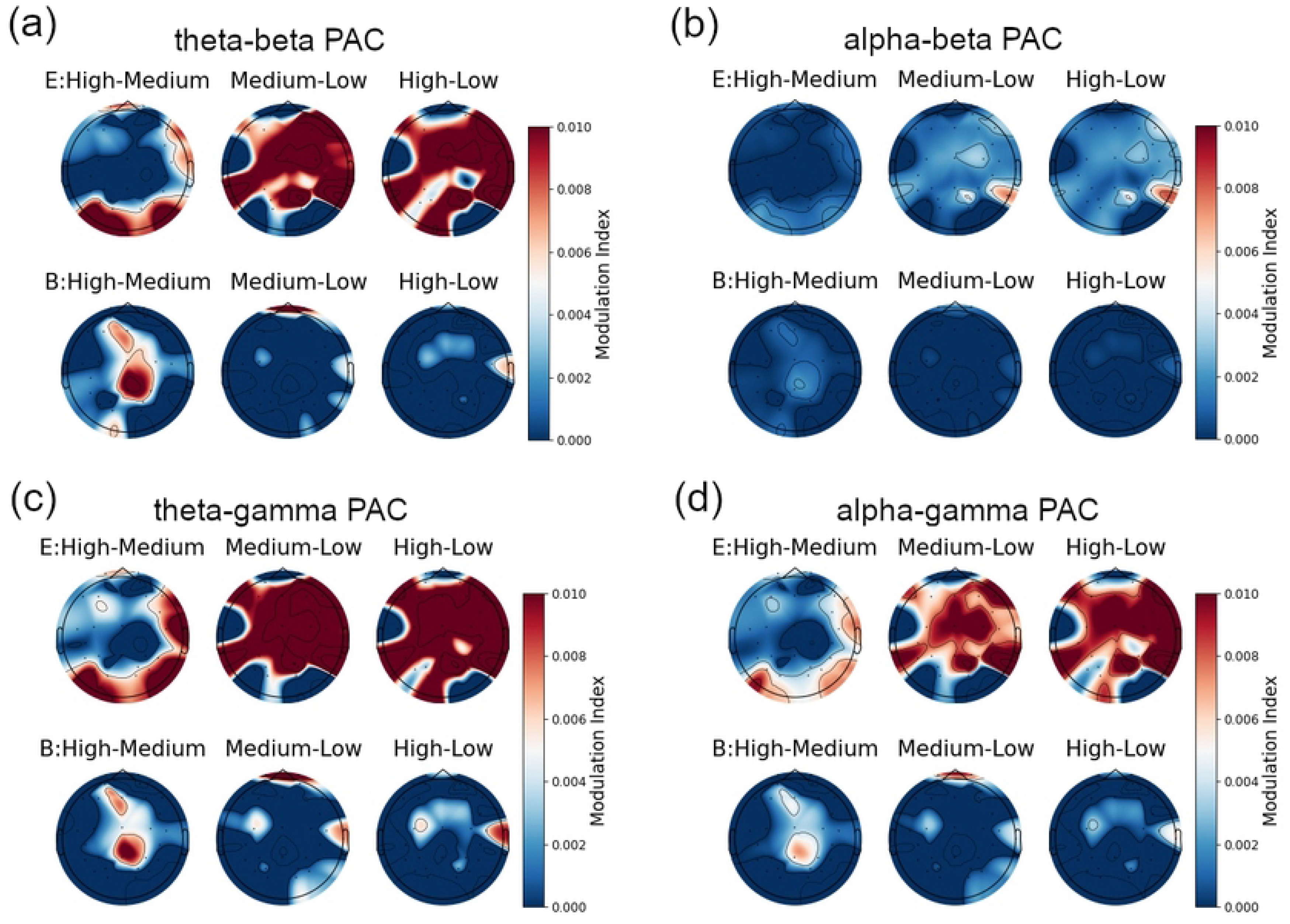
Topography maps showing MI changes with cognitive load levels variations in simultaneous interpretation. (a) theta-beta PAC. (b) alpha-beta PAC. (c) theta-gamma PAC. (d) alpha-gamma PAC. E: Experienced interpreters. B: Beginner interpreters. Graphs were adjusted so that the maximum MI value, shared by all PACs, is 0.010.

## Discussion

The results of subjective and objective evaluations (biometric measurements by PAC) did not align between experienced participants and beginners (Figs 2 and 3). Furthermore, only the PAC analysis enabled the evaluation of the interaction between years of work experience and cognitive load level (Table 1 and Fig 2). The reason appears to be that subjective evaluations were conducted after the EEG measurements, focusing solely on the familiarity of the words from a subjective perspective. Conversely, the PAC analysis encompasses the entire process that occurred during SI. These results imply that PAC analysis, based on objective biological information, offers substantial insights into the cognitive load experienced during SI, beyond what is captured by subjective evaluations.

Based on the results of this study, we observed an increase in the theta-gamma PACs during SI among the experienced, and this increase was associated with an increase in the cognitive load (Table 3 and Fig 3 (c)). These findings emphasize the crucial roles played by the theta and gamma frequency bands in cognitive processes. Theta oscillations integrate new information and consolidate memories, while gamma oscillations coordinate information across different brain regions and process complex cognitive tasks, as reported in previous studies [37, 38]. Therefore, the coupling of theta and gamma bands is particularly significant in SI, where rapid language-switching and immediate information integration are required. Furthermore, the theta-gamma PAC has been reported to regulate the timing and synchronization of neural activity [33, 41]. Additionally, an increase in the cognitive load of working memory is associated with enhanced theta-gamma PAC activity, which has been reported to support effective stimulus processing and the conveyance of critical information related to perceptual and mnemonic representations [43]. This suggests that in high-speed or complex interpretation scenarios, the enhancement of theta-gamma PAC activity could aid maintaining focus and processing speed during SI. Such heightened activity might assist experienced interpretaters in efficiently performing under demanding conditions, ensuring accurate and fluent interpretations. Overall, combining previous research with these current results suggests the significance of theta-gamma PAC in the cognitive processes involved in SI, particularly in scenarios with high demands.

**Table 3.** Results of kruskal-wallis and dunn’s tests within cognitive load levels for experienced and beginner. L: low, M: medium, H: high : *p-value*: *p <* 0.05^*^, *p <* 0.01^**^, *p <*0.001^***^

| PAC types | Work experience | H Statistic | Kruskal’s p-value | $\eta_p^2$ | Load 1 vs. Load 2 | Dunn’s p-value |
| --- | --- | --- | --- | --- | --- | --- |
| Theta-beta PAC | Experienced | 18.0 | *** | 0.018 | $L < M$ | *** |
| Theta-beta PAC | Experienced | | | | $M < H$ | 0.755 |
| Theta-beta PAC | Experienced | | | | $L < H$ | *** |
| Theta-gamma PAC | Experienced | 16.3 | *** | 0.016 | $L < M$ | 0.003** |
| Theta-gamma PAC | Experienced | | | | $M < H$ | 0.279 |
| Theta-gamma PAC | Experienced | | | | $L < H$ | *** |
| Alpha-beta PAC | Experienced | 19.6 | *** | 0.019 | $L < M$ | *** |
| Alpha-beta PAC | Experienced | | | | $M < H$ | 0.670 |
| Alpha-beta PAC | Experienced | | | | $L < H$ | *** |
| Alpha-gamma PAC | Experienced | 16.6 | *** | 0.016 | $L < M$ | 0.004** |
| Alpha-gamma PAC | Experienced | | | | $M < H$ | 0.251 |
| Alpha-gamma PAC | Experienced | | | | $L < H$ | *** |
| Theta-beta PAC | Beginner | 6.70 | 0.035* | 0.006 | $L > M$ | 0.016* |
| Theta-beta PAC | Beginner | | | | $M < H$ | 0.658 |
| Theta-beta PAC | Beginner | | | | $L < H$ | 0.043* |
| Theta-gamma PAC | Beginner | 1.60 | 0.568 | 0.001 | $L < M$ | 0.231 |
| Theta-gamma PAC | Beginner | | | | $M < H$ | 0.818 |
| Theta-gamma PAC | Beginner | | | | $L < H$ | 0.325 |
| Alpha-beta PAC | Beginner | 6.21 | 0.044* | 0.006 | $L > M$ | 0.019* |
| Alpha-beta PAC | Beginner | | | | $M < H$ | 0.622 |
| Alpha-beta PAC | Beginner | | | | $L < H$ | 0.057 |
| Alpha-gamma PAC | Beginner | 1.79 | 0.516 | 0.008 | $L < M$ | 0.179 |
| Alpha-gamma PAC | Beginner | | | | $M < H$ | 0.921 |
| Alpha-gamma PAC | Beginner | | | | $M < H$ | 0.333 |

In our study, we observed an increase in alpha-gamma PACs among experienced interpreters as their cognitive load increased during SI (refer to Table 3 and Fig 3 (d)). The coupling between alpha and gamma frequency bands is particularly important for adapting brainwave activity based on the content of working memory and the nature of the task. Such adaptability becomes even more evident in delayed match-to-sample tasks in which participants temporarily remember a presented stimulus and later compare it with a subsequent stimulus to determine a match [55, 56]. This process requires active maintenance of working memory and the instantaneous processing of information. Studies involving visual-delayed match-to-sample tasks have shown that changes in working memory load fuel an increase in the alpha-gamma PAC activity in both the frontal and occipital lobes [57]. Perhaps the brain dynamics during SI are similar. Interpreters must immediately comprehend and translate spoken words, a task that demands exceptional attention, memory retention, and language processing abilities during SI. Alpha bands help focusing on relevant information by reducing irrelevant details, and gamma oscillations enable the synchronization of information across different brain regions, supporting complex cognitive processing. Therefore, the observed increase in alpha-gamma PACs among experienced interpreters during SI likely signifies their brain’s efficient adaptation to task demands.

Based on our findings, we observed that as cognitive load increases during SI, the MI values of alpha-beta PACs increase (Table 3 and Fig 3 (b)). These findings align with previous research which concluded that alpha bands inhibit the processing of unnecessary information, while beta bands have been associated with maintaining network configurations, supporting schema retention during language transitions [69, 70]. In SI, interpreters frequently switched between their native language (Japanese: L1) and the target language (English: L2). This step required suppressing L1 due to differences in word order between Japanese and English, resulting in a high cognitive load for information maintenance [4]. In Japanese, the object is usually placed before the subject. Therefore, when translating from Japanese to English, it is necessary to retain the information of the Japanese object in memory until the term appears in the English translation. Furthermore, it is considered that the cognitive demands for retaining information increase when dealing with unfamiliar vocabulary. Our findings suggest that the interaction between alpha and beta PACs supports language selection during SI and network maintenance in high cognitive load on SI [71]. Our study aligns with these findings, showing an increase in alpha-beta PACs with higher cognitive load [62]. In SI, interpreters must understand the spoken content while maintaining the translated text in their working memory. The increase in the context of the highly complex task of SI, suggests that the beta band plays a role in regulating the strength of PACs in accordance with the complexity of the task, adapting to the demands of the task.

The MI values for the alpha-beta PAC were significantly lower than the other PACs, although they had a large effect size for interaction between cognitive load levels and work experience (Table 1). Perhaps the information transfer between the alpha and beta bands is limited. However, the significant effect size indicates that this limited information exchange is important for cognitive adjustments. Griffiths et al. and Park et al. argued that a decrease in the alpha-beta oscillations in the occipital region, coupled with an increase in the theta-gamma PAC in the hippocampus, supports memory consolidation [12, 44, 61]. Furthermore, Hanslmayr and colleagues emphasized the importance of a decrease in the alpha and beta oscillatory power for encoding episodic memories into the long-term memory [72]. From an information theory perspective, this decrease in alpha-beta oscillations indicates unpredictable states, which may convey essential information in information transfers [72]. Considering these previous studies, the decrease in alpha-beta oscillations may provide critical information as the brain processes new information and encodes it into long-term memory. Especially in complex tasks like SI, which demand sophisticated cognitive functions for focusing on necessary information from episodic memory, the decrease in beta oscillations could reduce the MI values.

Our analysis revealed an increase in theta-beta PAC when cognitive load increased during SI, which aligns with prior research conducted by Doume et al [58](Fig 3(a)). According to their findings, maintaining sensory information in working memory is connected to strengthening theta-beta PAC. Their studies suggest that enhanced phase synchronization from the medial temporal lobe area to the lateral prefrontal cortex in audio-visual working memory tasks indicates the impact of high cognitive load tasks, such as SI, on brain communication mechanisms, including theta-beta PAC and inter-regional phase synchronization. These results imply that SI requires enhanced communication between brain regions for cognitive control and the integration of multisensory information. Our study observed a significant difference in theta-beta PAC across a broad range of areas, including the frontal lobe, as cognitive load increased during interpretation (Fig 4 (a)). Our results suggest how complex cognitive tasks such as SI likely impact the brain’s communication mechanisms to maintain working memory.

Our findings indicate that interpreters with varying levels of experience exhibit distinct patterns of brain activity in response to cognitive load. Previous research by Yokota et al. demonstrated that as the cognitive load of the N-back task increased, the auditory steady-state response (ASSR) to 40-Hz pulses decreased, suggesting limitations in memory resources [73]. In contrast, our study found that experienced participants exhibited an increase in PAC under cognitive load, unaffected by memory resources. The presence of the so-called “interpreter’s advantage,” observed at the behavioral level in experienced interpreters, may result from enhanced cognitive abilities due to exposure to the demanding multitasking requirements of SI [1]. Additionally, research by Dimitriadis et al. indicates that as tasks become more complex, the strength of the beta-band PAC decreases [60]. In the beginner interpreters in our study, the only observed decrease in PAC with increased load was in the theta-beta and alpha-beta PACs, which include beta bands (Figs 3 (a) and (b)). Perhaps beginners require different training approaches to manage cognitive load, whereas experienced interpreters may benefit from various techniques to enhance their performances.

Our studies suggest that experienced interpreters demonstrate extensive neural activity across numerous brain regions (Fig 4). This suggests that experienced interpreters have simultaneously developed extensive brain regions through years of training and practice, enabling them to efficiently organize a variety of processes, such as listening, understanding, memory retention, and language production. Beginners have not yet sufficiently developed such specialized or adapted brain areas, which is likely why no increase was observed in response sites associated with cognitive load. Previous research suggests that changes in working memory load influence the interactions between various brain regions. Specifically, studies employing tasks with varying cognitive demands conclude that working memory load impacts beta-band activity in the right hemisphere, a crucial location for coordinating frontal control with sensory processing in complex tasks [65]. The interaction between the brain’s frontal and posterior regions in the beta-band changes based on the working memory load, emphasizing its importance in the brain’s regulatory functions. An increase in theta-beta PAC during the delay period of working memory tasks has also been reported, associated with the synchronization of different brain regions, thereby enhancing cognitive processing [58].

In the context of SI, such neuroscientific mechanisms become particularly crucial. SI demands high-level listening, immediate comprehension, accurate memory retention, and swift language production, all of which significantly increase working memory load. Experienced interpreters manage this cognitive load by developing specific brain regions and through robust interactions between different brain areas. These neuroscientific interactions enhance the cognitive processing necessary for simultaneous interpreters to process multiple streams of linguistic information and produce accurate translations.

## Conclusions

This study clarified that experienced interpreters demonstrate various PAC patterns across multiple frequency bands, which increase as cognitive load increases. In contrast, beginner interpreters showed no significant increase. Despite the relatively small MI values of the alpha-beta PAC effect, the observed effect size in the interaction between interpreters’ years of experience and cognitive load was large. This finding indicates that alpha-beta PAC may play a critical role in the relationship between interpreters’ years of experience and cognitive load.

Experienced interpreters exhibited a positive moderate correlation between cognitive load and PAC, whereas beginners displayed a weak negative correlation. These findings suggest that interpreters adopt unique strategies to manage diverse cognitive loads, and experienced interpreters likely possess well-developed neural processing strategies to handle the demands of complex information integration and attention control.

However, we must acknowledge the limitations of this study, including the uneven distribution of subjects between experienced interpreters (7) and beginners (15) and the absence of comparisons between first language (L1) and second language (L2). Future research will explore inter-professional comparisons related to cognitive load and measure the effectiveness, particularly within various fields of application.

